# Sequence specificity in DNA binding is determined by association rather than dissociation

**DOI:** 10.1101/2021.04.08.438955

**Authors:** Emil Marklund, Guanzhong Mao, Sebastian Deindl, Johan Elf

## Abstract

Sequence-specific binding of proteins to DNA is essential for accessing genetic information. Here, we derive a simple equation for target-site recognition, which uncovers a previously unrecognized coupling between the macroscopic association and dissociation rates of the searching protein. Importantly, this relationship makes it possible to recover the relevant microscopic rates from experimentally determined macroscopic ones. We directly test the equation by observing the binding and unbinding of individual *lac* repressor (LacI) molecules during target search. We find that LacI dissociates from different target sequences with essentially identical microscopic dissociation rates. Instead, sequence specificity is determined by the efficiency with which the protein recognizes different targets, effectively reducing its risk of being retained on a non-target sequence. Our theoretical framework also accounts for the coupling between off-target binding and unbinding of the catalytically inactive Cas9 (dCas9), showing that the binding pathway can be obtained from macroscopic data.

**One Sentence Summary:** Association and dissociation rates are anti-correlated for reactions that include a nonspecific probing step.

## Main Text

Sequence-specific recognition and binding of DNA target sites by proteins such as polymerases, DNA-modifying enzymes, and transcription factors are essential for gene expression and regulation across all kingdoms of life (*1*). The textbook explanation for this sequence dependence of binding posits that favorable hydrogen bonding interactions between the protein and particular DNA sequences result in prolonged binding times (*2*). Consequently, the rate of protein dissociation would depend on the DNA sequence, while the association rate would be invariant with respect to sequence. Indeed, the rate of protein association with DNA has often been assumed to be sequence-independent (*3–6*). However, single-molecule measurements have shown that when a protein scans the DNA for binding sites, the association rate does depend on the sequence (*7*), and that different target sequences can be bypassed with distinct probabilities (*8*). These differences have been ascribed to differences in the probability of recognition when the protein is centered on the target sequence (*7*). It is unknown whether the rate of binding imposes constraints on the rate of dissociation, beyond the fact that the ratio of association and dissociation rates is necessarily dictated by the free energy difference between the free and bound states. So far, the recovery of microscopic parameters from experimentally obtained macroscopic ones has remained an unsolved problem, limiting our understanding of how sequence-specific binding is achieved on the microscopic level.

To explore the limits of the association and dissociation rates, we considered the standard model (*9*), according to which a protein has a non-specific testing mode where it is bound nonspecifically to DNA (Fig. 1A). In the testing state, the protein can either specifically bind the target with probability *p*_tot_, or dissociate into solution with probability 1-*p*_tot_. When the association process is modeled as a three-state (specifically bound, nonspecifically bound, and dissociated) continuous time Markov chain, the effective macroscopic target association and dissociation rates (*k*_a_ and *k*_d_) relate to each other according to (see Supplementary Text for derivation)

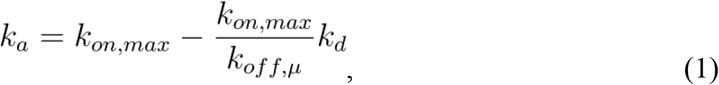

where *k*_on,max_ is the association rate given by a searching protein that binds the target upon every non-specific encounter (*p*_tot_=1), and *k*_off,μ_ is the rate of microscopical dissociation from the bound state into the nonspecifically bound searching mode. This equation implies that the association and dissociation rates are inherently coupled, and linearly anti-correlated if binding sites exhibit identical microscopical dissociation rates, since *k*_on,max_ does not depend on the specific sequence. The linear relationship between *k*_a_ and *k*_d_ described by Eq. 1 is implicitly parameterized by the probability of binding rather than dissociating from the non-specifically bound state, *p*_tot_, such that an increase in *p*_tot_ causes an increase in *k*_a_, and a corresponding decrease in *k*_d_ (Fig. 1B). This anticorrelation can be intuitively understood by acknowledging that a decrease in the number of target site encounters required for successful binding must, in turn, result in a corresponding increase in the number of dissociation attempts needed for macroscopic dissociation from the target (Fig. 1C). Most importantly, Eq. 1 makes it possible to obtain a microscopic parameter, such as *k*_off,μ_, from macroscopically measurable parameters, such as *k*_a_ and *k*_d_.

**Fig. 1.**
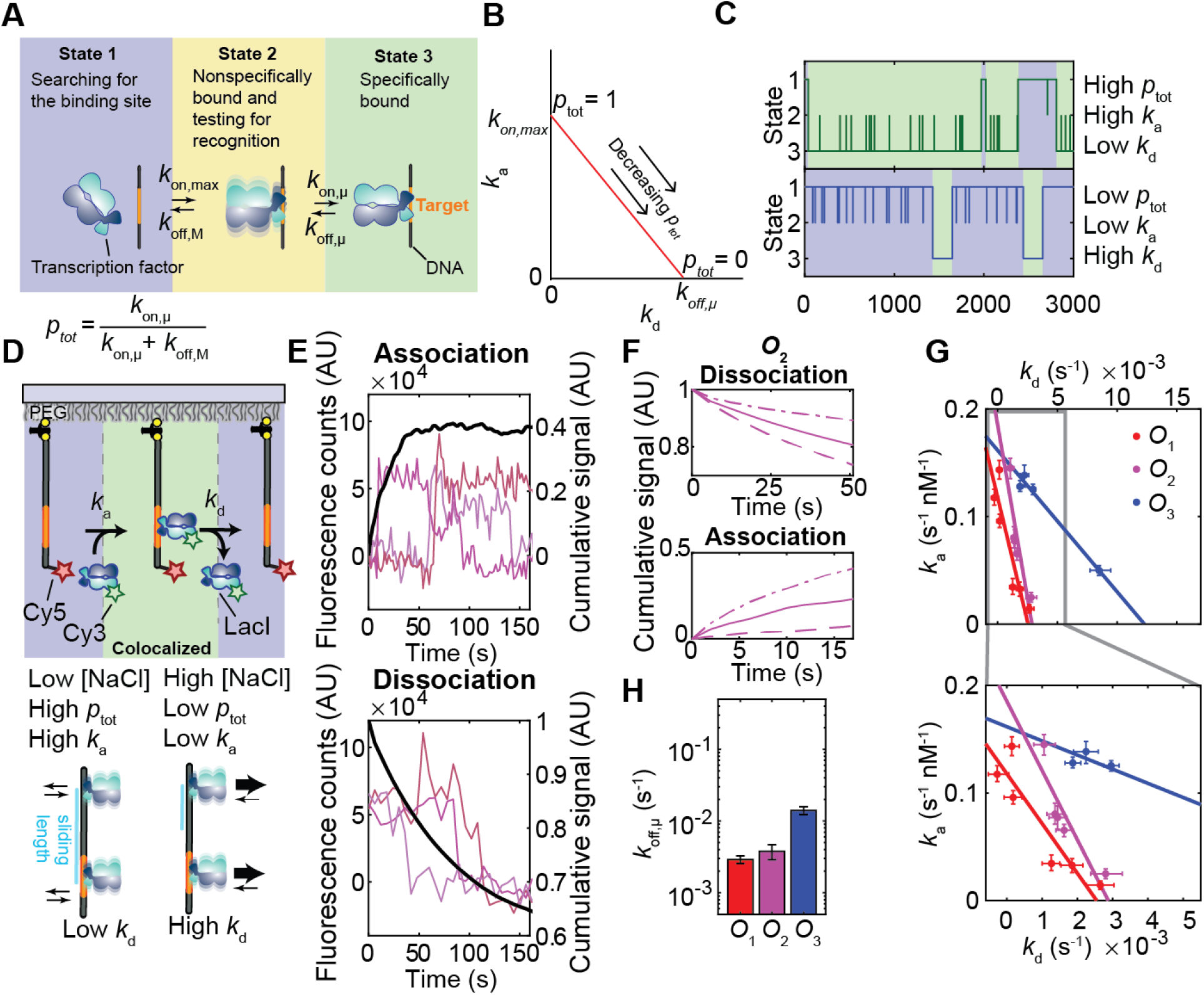
Bimolecular association and dissociation rates are inherently anti-correlated due to target-site probing. (**A**) Schematic of the kinetic model describing protein-DNA binding. (**B**) The effective rate constants for the association to (*k*_a_) and dissociation from (*k*_d_) the target site are coupled according to Eq. 1. This relationship becomes anti-correlated and linear when *k*_off,μ_ is constant and *p*_tot_ changes (red line). (**C**) Example traces from stochastic simulations sampling the association, dissociation, and nonspecific binding with target-site probing. When *p*_tot_ is high (top), the search times become short (1/*k*_a_, blue areas) and the binding time long (1/*k*_d_, green areas). When *p*_tot_ is low (bottom), the search times become long and the binding times short. (**D**) Single-molecule colocalization measurements detect association and dissociation for LacI binding to its operators (left) and the predicted effect on association and dissociation rates of changing the salt concentration (right). (**E**) Example single-molecule traces showing binding to and unbinding from the *O*_2_ operator at 100 mM NaCl (colored lines) and the normalized association and dissociation curves (black lines) obtained after summing 667 and 773 traces for the association and dissociation experiment, respectively. a.u., arbitrary units. (**F**) Normalized association and dissociation curves for *O*_2_ and 1 mM (dashed dotted), 100 mM (solid), and 200 mM NaCl (dashed). (**G**) Measured *k*_a_ and *k*_d_ values for the three *lac* operators, and fits to Eq. 1 for each repeat (colored lines). The salt concentrations used for the different experiments are in the range 1-250 mM supplemented NaCl for *O*_1_ and *O*_2_, and 1-100 mM supplemented NaCl for *O*_3_ (Fig. S1). Error bars are 68% confidence intervals obtained after bootstrapping the fluorescence traces in each experiment. (**H)** Microscopic dissociation rates *k*_off,μ_ for the different operators, estimated as the *k*_d_-intercepts of the fits to Eq. 1. Error bars are standard errors from *n* = 2 salt titrations for each operator.

To experimentally test the anti-correlation between association and dissociation rates, we measured the kinetics with which a prototypical DNA-binding protein, the transcription factor LacI, binds to its natural operator sites using single-molecule fluorescence colocalization. We surface-immobilized a Cy5-labeled DNA construct containing a natural *lacO* operator site (*O*_1_, *O*_2_, or *O*_3_) and used total-internal-reflection fluorescence microscopy to monitor individual DNA molecules (Fig. 1D). Upon addition of LacI labeled with Cy3 distal from the DNA binding domain, we monitored the appearance and disappearance of well-defined spots with co-localized fluorescence emission from both Cy3 and Cy5 (Fig. 1E). The Cy3 label has previously been shown to affect neither the specific nor the nonspecific DNA binding (*8*) (Labeling efficiency: 84.5 %; see also Supplementary Text and Table S1). Few DNA molecules featured co-localized LacI-Cy3 spots in control experiments with Cy5-labeled DNA constructs lacking an operator site (11% and 3% at 1 and 100 mM NaCl, in contrast to >65%, >60% and >20% at 1, 100 and 200 mM NaCl for DNA with an *O*_1_ operator; Fig. S1), indicating that the Cy3 spots represent complexes of LacI-Cy3 specifically bound to the operator with only a minor contribution from nonspecific binding of LacI-Cy3 to DNA or to the surface.

To implement conditions that give rise to a range of different association and dissociation rates, we varied the salt concentration in our experiments (Fig. 1F) since changes in salt concentration are expected to affect the time that LacI spends nonspecifically bound to DNA while sliding along it (*9*, *10*). This in turn would change the number of operator encounters per nonspecific association, such that *p*_tot_ is expected to increase with decreasing salt concentration. We note that the *k*_a_ values measured for each salt titration should be interpreted as being merely proportional to the true bimolecular association rate constants since the exact concentration of active LacI needed for normalization can vary between salt titration repeats due to differences in the extent of protein surface adsorption, protein stability, and pipetting errors. Nevertheless, we obtain a reproducible and anti-correlated relationship between the measured *k*_a_ and *k*_d_ values for each salt titration and operator, consistent with the notion that *p*_tot_ varies, while *k*_off,μ_ remains constant for each operator when changing the salt concentration (Fig. 1G; see also Fig. S2–4).

To test if the specificity of LacI-binding to different operators is due to differences in microscopic association or dissociation, we fit Eq. 1 to the experimentally determined *k*_a_ and *k*_d_ values (colored lines in Fig. 1G), yielding *k*_off,μ_ for each operator as the *k*_d_-intercept of each *k*_a_ *versus k*_d_ line (Fig. 1H). Surprisingly, the estimates obtained for *k*_off,μ_ are very similar for all operators.

Analogously, we can also estimate *k*_off,μ_ for the different operators (Fig. 2A) from existing *in vivo* estimates of *k*_a_ and *k*_d_ ((*7, 11–13*), Fig. 2B). *k*_on,max_ has been measured independently *in vivo* (*7*), and *k*_off,μ_ can therefore be calculated as the only unknown in Eq. 1. Consistent with what we found in our *in vitro* experiments, the *k*_off,μ_ estimates obtained from *in vivo* data are very similar for all operators (Fig. 2C). Even though the *K*_D_ value of *O*_2_ exceeds that of *O*_1_ more than 4-fold, and that of *O*_sym_ 20-fold, these operators exhibit essentially the same *k*_off,μ_ *in vivo*, as they all fall on the same *k*_a_ *versus k*_d_ line in Fig. 2B. The differences in *K*_D_ observed for the different operators can thus be explained predominantly by differences in target-site recognition (*p*_tot_) and the numerous microscopic re-associations that the protein undergoes before every successful macroscopic dissociation from a strong operator.

**Fig. 2.**
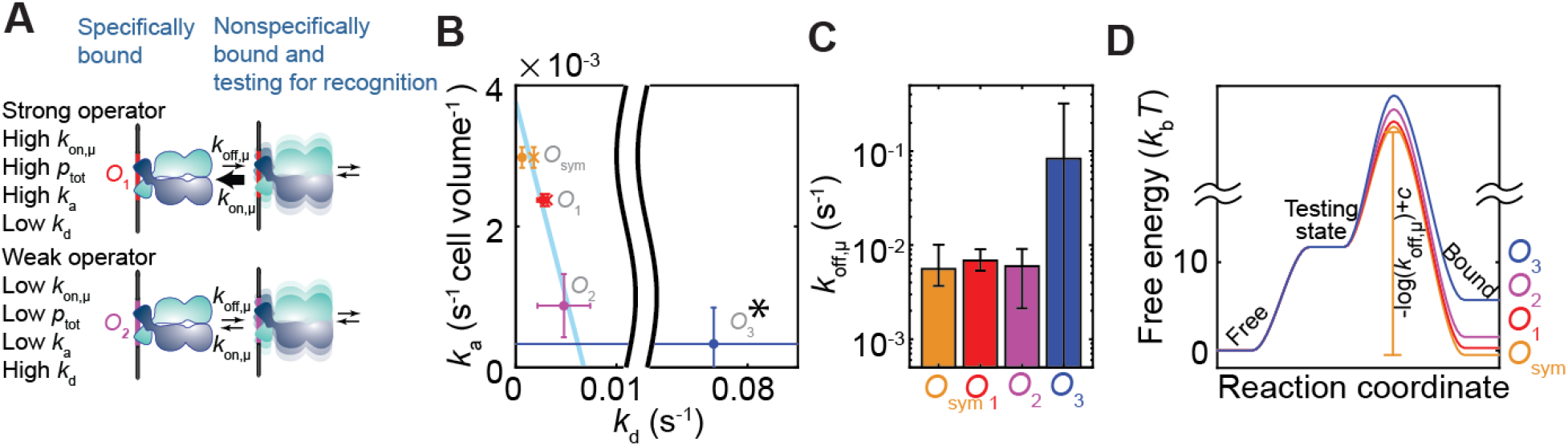
Different DNA targets exhibit similar microscopic dissociation rates but different recognition probabilities. (**A**) Predicted effect on the association and dissociation rates if *k*_on,μ_ were different but *k*_off,μ_ identical for the different operators. (**B**) Experimental single-molecule target-site association rates *k*_a_ (*7*) plotted against the dissociation rates *k*_d_ for the different *lac* operators. For the crosses, *k*_d_ was directly measured by single-molecule imaging (*15*). For the dots, *k*_d_ was calculated as *K*_D_×*k*_a_, where the equilibrium constant *K*_D_ was measured via the repression ratio of gene expression (*11*, *12*). Cyan line, best fit of Eq. 1 to the *O*_sym_, *O*_1_ and *O*_2_ data. * Due to the large error in the *k*_d_ estimate for *O*_3_ (68% CI: [−0.04,0.20] s^−1^), it has been excluded from the fit. Error bars are standard errors, obtained by propagating the errors from the experiments. (**C**) Microscopic dissociation rates *k*_off,μ_ for the different operators, estimated from the *in vivo* data using Eq. 1. Error bars are 68 % confidence intervals, obtained by propagating the errors from the experiments. The confidence interval for is *O*_3_ is [−0.05,0.32] s^−1^, making this *k*_off,μ_ estimate an upper bound. (**D**) Energy landscapes (a putative rather than true reaction coordinate is shown) for the transition from free (State 1) to bound (State 3) states for the different operators, as determined by the measured *K*_D_ and *k*_off,μ_ values. The activation energy on the transition path between the testing state and bound state is not uniquely determined, but the differences in activation energies between the different operators are. The activation energy is equal to −log(*k*_off,μ_) + *c*, where *c* is the same constant for all operators (see Supplementary Text).

The agreement between the experimental data and our simple model suggests that LacI binding dynamics can be captured with one kinetic barrier, the height of which differs for different operators; i.e., it is more favorable for LacI to bind to certain operators than to others when sliding by, but the rate of escaping from the specifically bound state does not depend on the sequence (Fig. 2D). By recognizing that mutations along the binding pathway can be seen as energetic barriers for binding, our theoretical framework can also be used to dissect the binding path in more complex, sequential binding mechanisms. Accordingly, one would first mutate a binding sequence in several different ways, measure the resulting macroscopic rates *k*_a_ and *k*_d_, and then determine which sector of the (*k*_a_,*k*_d_)-space the different mutations fall into (Fig 3A). Assuming that the native sequence has the highest *k*_a_ value and that mutations introduce a rate-limiting step, the sectors will be ordered according to the position of the mutations along the reaction pathway. Thus, ratelimiting steps closer to the bound state will result in fewer rebinding events, leading to an increase in *k*_d_ for the same value of *k*_a_.

**Fig. 3.**
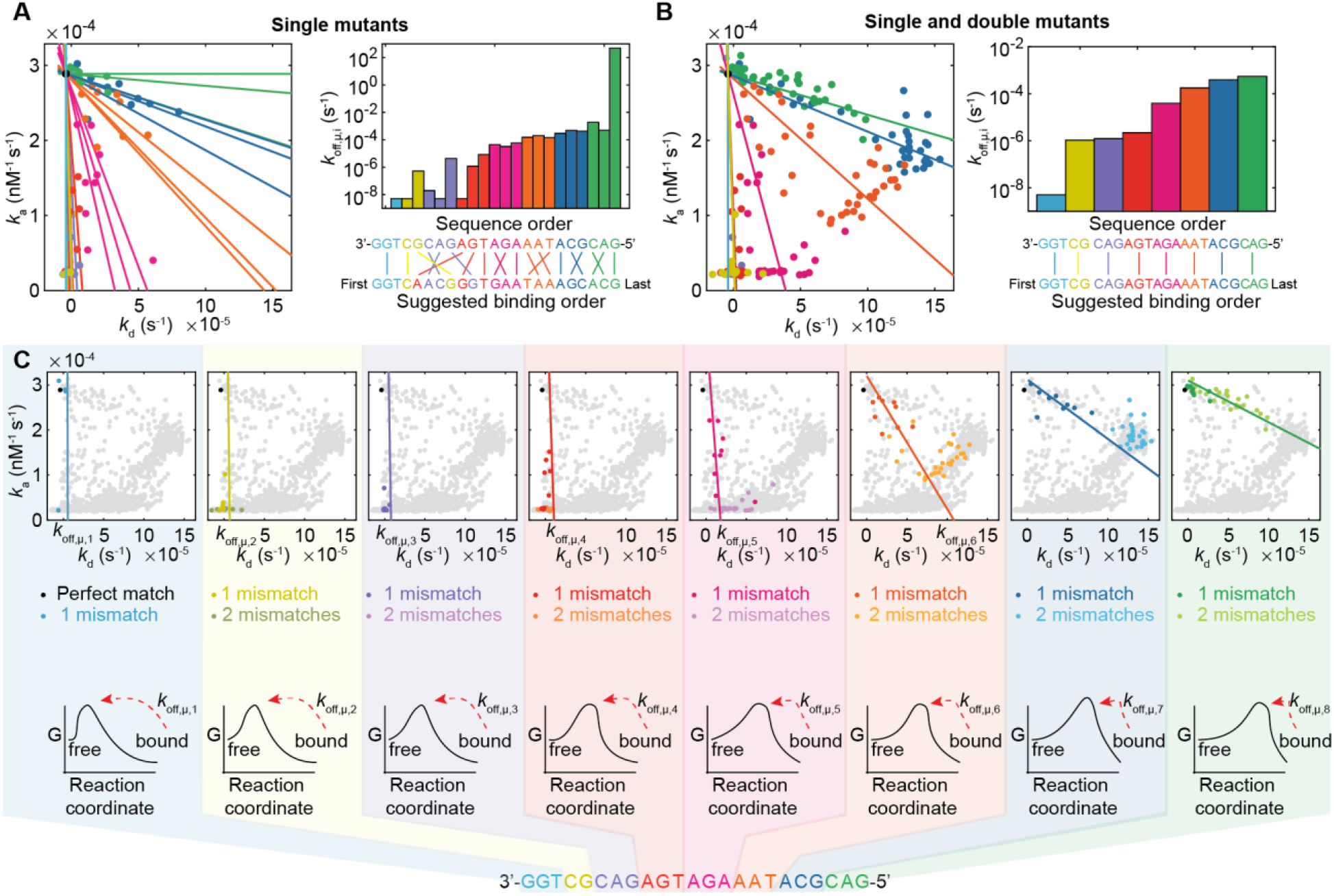
The target-site recognition model describes the coupling between dCas9 off-target binding and unbinding. Triplets along the sequence are indicated by different colors. (**A**) Measured association and dissociation rates for dCas9 and different off-target sequences (single-site mutant data from (*14*)) (left). Linear fits to the data for each mutated base yield the estimated effective microscopic dissociation rate *k*_off,μ,i_ (i.e. *k*_d_-intercept) associated with each nucleotide, indicating the binding pathway (right). (**B**) Measured association and dissociation rates for dCas9 and different off-target sequences with single and double mismatches (*14*) (left), and estimated effective microscopic dissociation rate *k*_off,μ,i_ associated with each 3-nucleotide DNA region, indicating the binding pathway (right). (**C**) Measured association and dissociation rates for dCas9 and different off-target sequences (grey dots; single and double mismatch mutant data from (*14*)), where data for mismatches occuring in specific gRNA regions are highlighted in colors. In (**A**) and (**B**), colored lines are individual fits of Eq. 1 to the single, or the single and double mismatch data, respectively. In (**C**), colored lines are representations of a global fit of a model with an 8-state sequential recognition to all the data. Here each line shows the model-predicted effect of mutations in one specific gRNA triplet region (see Supplementary Text and Fig. S5). Note that some of the data points in the plots are not highlighted in colors. These data points represent sequences with mutations in two different gRNA regions. In the right panels of (**A**) and (**B**) estimated *k*_off,μ,i_ values smaller than 5 × 10^−9^ s^−1^ have been rounded up to this value.

To demonstrate the use of this method, we apply it to high-throughput association and dissociation data available for dCas9 binding to off-target, mismatch mutants ((*14*), Fig. 3). dCas9 is guided by an RNA (gRNA) when it binds DNA, with a reaction coordinate for the testing of recognition that is already well-established (*15*-*17*). The gRNA binds the DNA by base pairing in a sequential manner starting from a seed sequence, and then continuing hybridization at base pairs more distal from the seed. When we plot linear fits to the single-base mismatch data, the resulting slopes and *k*_d_-intercepts indicate a binding pathway that is well-aligned with the order of basepairs in the guide RNA sequence, starting at the seed and then moving further into the sequence (Fig. 3A). When we group the sequence into 3-nucleotide groups and plot the lines corresponding to single- and double-base mismatches within each group, the *k*_d_-intercepts show a binding pathway that is identical to the order of the base pairs in the guide RNA sequence (Fig 3B). Based on these groups, we have colored all the single and double mismatch mutations in the *k*_a_ *versus k*_d_ plots of Fig. 3. The resulting rainbow-colored pattern, the order of which corresponds to the order in the guide RNA sequence, demonstrates how the *k*_a_ *versus k*_d_ plot maps the mutations onto the reaction coordinate (Fig 3C).

In conclusion, the efficiency of target-site recognition not only is crucial for determining protein-DNA association rates but also plays an equally important role in determining how long proteins remain bound to their targets. In the case of the *lac* repressor, we have shown that the efficiency of target-site recognition (*p*_tot_) - and *not* how long the protein remains in the bound state - causes the differences in binding strength observed for different sequences. This behavior may represent an evolutionary adaptation to facilitate fast search by minimizing the risk of the protein being retained on sequences that resemble the actual operators. The coupling between association and dissociation rates holds for all bimolecular association-dissociation processes adhering to detailed balance, where a step of rapid testing for molecular recognition precedes the strong binding of a target. Our theoretical result is therefore very likely to be generally applicable to a wide range of kinetic systems in addition to the ones investigated here, including processes that do not involve protein-DNA interactions.

## Acknowledgments

We thank Otto Berg, Måns Ehrenberg, Jakub Wiktor, David Fange, Irmeli Barkefors, and Daniel Jones for discussions.

## Author contributions

JE and EM conceived the study; EM derived Eq. 1; SD, EM, and MG designed the experiments; MG performed the experiments; EM analyzed the data; EM, SD and JE interpreted results; EM, JE, and SD wrote the paper;

## Data and materials availability

All raw data and analysis codes will be made available upon request.

## Supplementary Materials for

### Material and Methods

#### Expression, purification and fluorophore labelling of the *lac* repressor

Cy3 labelled *lac* repressor dimer (LacI-Far-2) was prepared according to previously published methods (*8, 18*), with a Cy3 introduced distal from the DNA binding domain of LacI. Briefly, the protein contains a C-terminal 6xHis-tag for affinity purification purposes, and the C-terminal tetramerization domain has been removed. A cysteine for labeling was introduced at amino acid position 312. All cysteines found in the wild-type protein were removed from the sequence, except for the solvent-excluded cysteine in the monomer-monomer interface required to maintain an intact dimer (*18*).

#### DNA constructs for single-molecule fluorescence co-localization measurements

Double-stranded DNA constructs that contained operator sites as indicated, a backbone-incorporated Cy5 fluorophore attached to position 5 of a dT base via a 6-carbon linker (Integrated DNA Technologies), and an end-positioned biotin moiety were generated by annealing and ligating a set of overlapping, complementary oligonucleotides. High-performance liquid chromatography (HPLC)-purified oligonucleotides were mixed at equimolar concentrations in 50 mM Tris pH 8.0, 100 mM KCl, 1 mM EDTA, annealed with a temperature ramp (95-3°C), ligated with T4 DNA ligase (New England Biolabs), and purified by polyacrylamide gel electrophoresis (PAGE). Successful ligation was confirmed by denaturing PAGE.

#### Single-molecule colocalization microscopy

Biotinylated and fluorophore-labeled DNA constructs were surface-anchored onto PEG-coated quartz microscope slides through biotin-streptavidin linkage (*19, 20*). Cy3 and Cy5 dyes were excited with 532 nm Nd:YAG and 638 nm diode lasers, respectively, and fluorescence emissions from the two fluorophores were detected using a custom-built prism-based TIRF microscope, filtered with ZET532NF (Chroma) and NF03-642E (Semrock) notch filters, spectrally separated by 635 nm (T635lpxr) and 760 nm (T760lpxr) dichroic mirrors (Chroma), and imaged onto the separate regions of an Andor iXon Ultra 888 electron multiplying charge-coupled device (EMCCD) camera. Imaging was carried out in imaging buffer containing 20 mM K2HPO4:KH2PO4 pH 7.4, 1 mM β-Mercaptoethanol, 0.05 mM EDTA, 100 μg/ml acetylated BSA (Promega), 10% (v/v) glycerol, 10% (w/v) glucose, 0.01% Tween 20 (v/v), 2 mM Trolox to reduce photoblinking of the dyes (Rasnik et al), an enzymatic oxygen scavenging system (composed of 800 μg/ml glucose oxidase and 50 μg/ml catalase), as well as 1-300 mM NaCl (as indicated). LacI was introduced by infusing the sample chamber with imaging buffer supplemented with 0.5 nM LacI using a syringe pump (Harvard Apparatus). During image acquisition, a laser exposure time of 1 s was used. A frame rate of 0.5 Hz was used when detecting association and measuring *k*_a_, except for one of the salt titration repeats for *O*_3_ (cyan crosses, Fig. S2) where a 1 Hz frame rate was used. Directly following the association experiment, a movie at 1/6 Hz was collected for detecting dissociation and measuring *k*_d_, except for one of the salt titration repeats for *O*_3_ (green crosses, Fig. S2) where the association movie at 1 Hz frame rate was used for estimating dissociation. For one of the *O*_1_ salt titrations (orange and brown crosses in Fig. S2) an additional movie at 1/12 Hz was captured directly after the 1/6 Hz movie. For this salt titration, *k*_d_ values were estimated individually from the 1/12 Hz (brown crosses, Fig. S2) and 1/6 Hz (orange crosses Fig. S2) movie, while the same 0.5 Hz movies were used for estimating the corresponding *k*_a_. All *k*_a_ and *k*_d_ were estimated from individual flow experiments except for the estimates for *O*_3_ at 15 mM and 35 mM NaCl. For these two data points *k*_a_ and *k*_d_ were estimated from two flow experiments each, captured directly after each other.

#### Analysis of single-molecule colocalization measurements

The data were analyzed by summing the Cy3 fluorescence intensities within a 7 x 7-pixel square for each frame and each Cy5 dot. An à trous wavelet decomposition was used for dot detection in the Cy5 images (*21*). Dots were detected through scale-dependent standard deviation thresholding in the second wavelet plane with a threshold of three standard deviations, where the standard deviation was estimated by the median absolute deviation method (*22*). Dot centers were localized by calculating the weighted centroid from the pixel regions obtained from dot detection. Background intensities for pixels were estimated by a 2D moving average of each Cy3-fluorescence image, with exclusion of outliers by assigning them lower weights when calculating the average (*23*), so as not to include pixels corresponding to fluorescent dots when calculating the moving average. The fluorescence counts for each trace and frame were calculated as the difference between the raw fluorescence signal and the local background in the Cy3 image. Cumulative fluorescence curves were obtained by aligning and summing regions of single-molecule traces that had a current startpoint of the region corresponding to either low (association) or high (dissociation) fluorescence (Fig 2E) values. For the association curves, the trace regions being aligned and summed over started in a three-frame window just after LacI was introduced into the flow channel, and ended at the end of the movie. For the dissociation curves, the trace regions being aligned and summed over started at any point in time more than 200 s before the end of the movie, and ended 200 s after the start point. Fluorescence traces were classified as ‘low fluorescence’ or ‘high fluorescence’ if the current count was below 20,000 or above 35,000, respectively. A threshold was set such that all counts above 50,000 were set to this value. Traces were excluded from the analysis if they had no counts higher than 50,000 within a 12-s moving-average window, indicating that no long-lived LacI binding occurred. To not bias results by including experiments with very weak binding, higher proportion of non-specific binding events compared to specific binding events, or higher proportion of binding to the glass surface, individual experiments were excluded from further analysis if they had a fraction of DNA spots with a binding event that was lower than 15% (See Fig. S1 for comparisons between operator and non-operator DNA). In total this excluded three data points at 200 and 250 mM NaCl from further analysis. For each experiment, the association and dissociation rates were estimated from the initial slopes of the cumulative curves. To account for photobleaching, we performed calibration dissociation experiments at different laser exposure times (fractional laser on times compared to the frame rate), and subtracted the constant contribution due to photobleaching from each *k*_d_ estimate (Fig. S3). The entire analysis pipeline was validated by performing stochastic simulations of binding and dissociation and simulated microscopy (*24*). This analysis method returned essentially the same association and dissociation rates as those that were put into the simulations (Fig. S4, Table S2). The simulations were performed with a number of DNA molecules that matches the average number of surface-immobilized DNA molecules found per field of view in the experiments (1400 Cy5 dots), and with imaging conditions mimicking the experiments in terms of level of background and fluorescence counts when LacI was bound. A frame rate of 0.5 Hz was used when simulating microscopy images for *k*_a_ estimation, and a frame rate of 1/6 Hz was used when simulating microscopy images for *k*_d_ estimation. The reported values for *k*_off,μ_ in Fig. 1H are mean ± standard error of mean (s.e.m.), where each sample was obtained by fitting Eq. 1 to data from individual salt titrations, and *n* = 2 independent titration experiments for each operator.

**Fig. S1.**
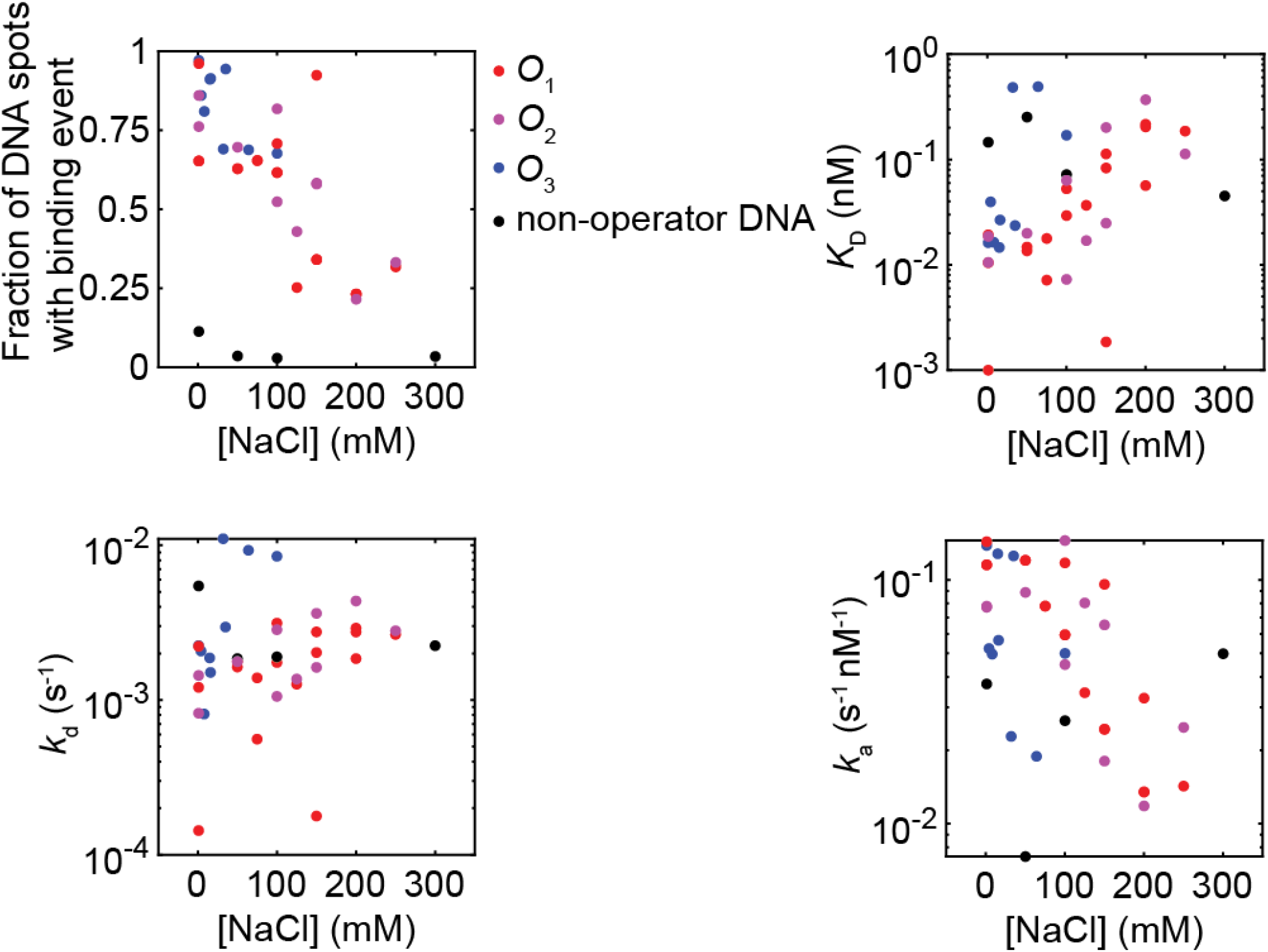
Binding and specificity in LacI-DNA colocalization measurements. The fraction of DNA spots that had at least one LacI binding event, as a function of the salt concentration of the experiment, for different operators and one non-operator DN (top left). A binding event was detected when a trace had a fluorescence count higher than 50,000 in a 12-s moving-average window. *K*_d_ (top right), *k*_d_ (bottom left) and *k*_a_ (bottom right) for the different DNA constructs as a function of the salt concentration of the experiment.

**Fig. S2.**
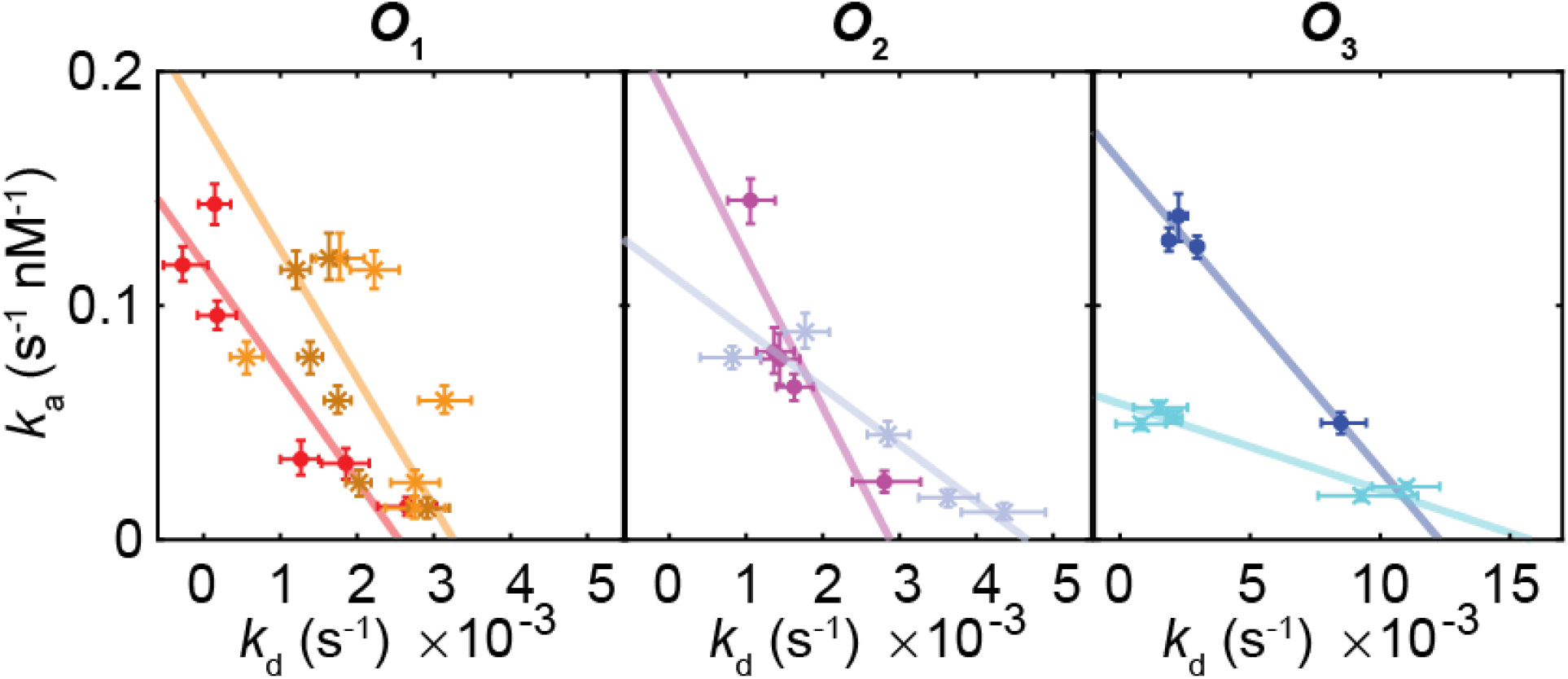
Single-molecule colocalization measurements for salt titration repeats. Measured *k*_a_ and *k*_d_ values from two salt titration repeats (dots and crosses) for the *lac* operators, and fits to Eq. 1 for each repeat (colored lines). For one of the *O*_1_ salt titrations, *k*_d_ was estimated with both 1/6 Hz (orange crosses) and 1/12 Hz (brown crosses) frame rate, while the corresponding *k*_a_ was estimated once with a 0.5 Hz frame rate. See Methods for additional experimental conditions. Error bars are 68 % confidence intervals obtained by bootstrapping the fluorescence traces.

**Fig. S3.**
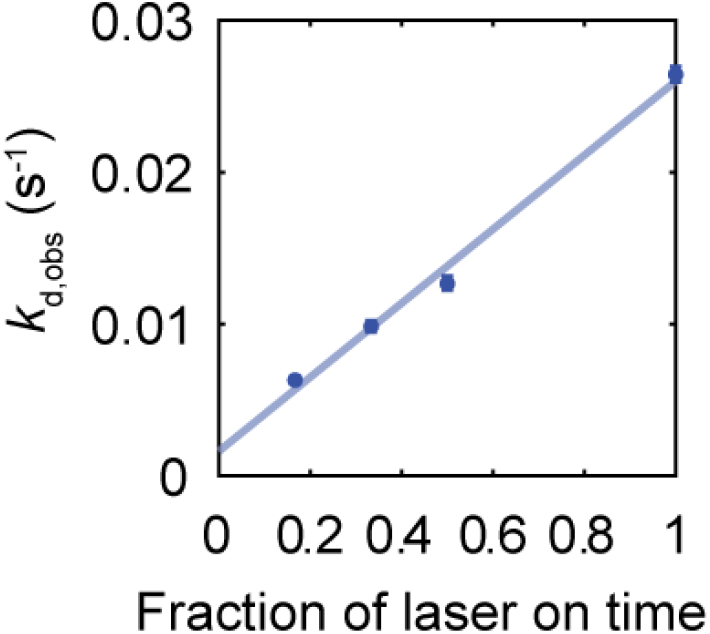
Bleaching calibration for *k*_d_ estimate. Observed dissociation rates *k*_d,obs_, obtained as the initial slopes of the dissociation curves, plotted against the fractional exposure time *f* used in the experiment. *k*_d,obs_ was measured for *O*_3_ dissociation at 1 mM NaCl, with a laser exposure time of 1 s, and frame rates of 1,0.5, 1/3 and 1/6 Hz (blue points). The blue line is the best fit to the equation *k*_d,obs_= *k*_d_+*f k*_bleach_, so that *f k*_bleach_ from the fit can be subtracted from *k*_d,obs_ to obtain *k*_d_ for each measurement. Error bars are 68 % confidence intervals obtained by bootstrapping the fluorescence traces in each experiment.

**Fig. S4.**
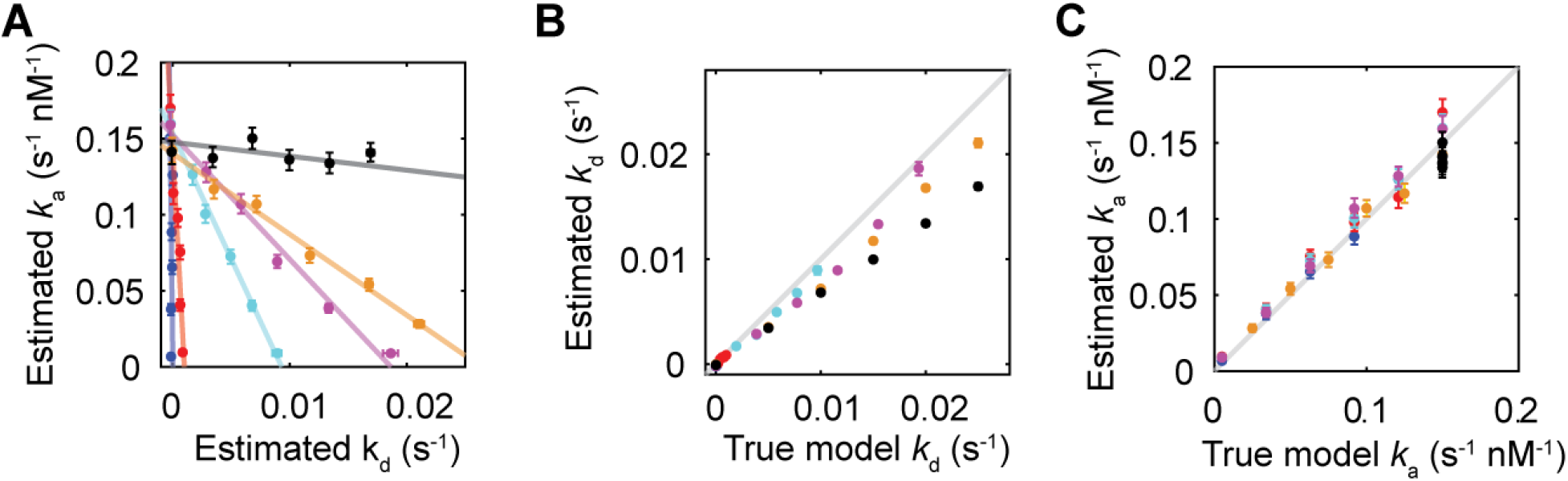
Stochastic simulations and simulated microscopy. (**A**) Estimated *k*_a_ and *k*_d_ for the simulated data. (**B**) True and estimated *k*_d_ values. (**C**) True and estimated *k*_a_ values. See Table S2 for true and estimated *k*_off,μ_ values for the simulated data. All error bars are 68 % confidence intervals obtained by bootstrapping the simulated fluorescence traces.

#### Analysis and regression of *in vivo* association and dissociation measurements

Measured values and associated errors for association rates (*7*), dissociation rates (*13*), and equilibrium constants (*11*, *12*) for *lac* repressor binding to different operators were obtained from their respective references. In (*11*, *12*) equilibrium data was measured as the fold-change of repression, and is reported in binding energies of the repressor to its operator. With the model used in (*12*), these binding energies can be recalculated to equilibrium constants via

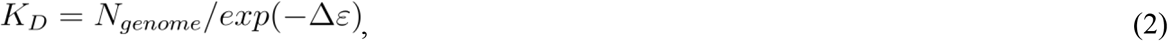

where *N*_genome_ = 5×10^6^ base pairs is the size of the *E. coli* genome, and *Δε* is the binding energy to the operator as defined in (*12*). In (*11*) and (*12*), the fold-change is measured for the LacI tetramer and dimer, respectively. The model fit of the data in (*11*) is shown to perfectly describe the data in (*12*), demonstrating that these constructs have identical binding energies. Thus, we can use the values of the binding energies and the associated errors from (*11*) when estimating *K*_D_ for the LacI dimer from Eq. 2. Association and dissociation rates are here reported in units per cell volume, and since ~ 4 *lac* repressor molecules were searching for the target sites in (*7*), all association rates were divided by 4. The association rates used here are taken from the additive model fit of all the data in (*7*), that is, they are extracted from the strains JE13, JE12, JE117, JE118, JE116, JE101 and JE104 containing different combinations of the *O*_sym_, *O*_1_, *O*_2_, and *O*_3_ operators. The reported values for *k*_off,μ_ were obtained by evaluating Eq. 1 with the experimentally estimated values for (*k*_a_,*k*_d_,*k*_on,,max_), where *k*_on,max_= *k*_a_/*p*_tot_, where *k*_a_ and *p*_tot_ are the values reported for *O*_1_ in (*7*). Error bars for *k*_off,μ_ are 68 % confidence intervals obtained by resampling (*k*_a_,*k*_d_, *k*_on,,max_) with the errors reported in (*7*, *11*-*13*), while assuming that the errors for all reported values are normally distributed.

#### Analysis and regression of dCas9 association and dissociation measurements

All data were taken from (*14*). Dissociation rates were taken from the dataset with chase, and association rates were obtained after combining the 1 nM and 10 nM datasets as described in (*14*). The colored lines in Fig. 3A and B were acquired by fitting Eq. 1 to the experimental data by minimizing the squared deviation between the model predicted (*k*_a_,*k*_d_) and the experimentally obtained (*k*_a_,*k*_d_), while constraining the line to go through the data point corresponding to the perfectly complementary sequence. In Fig. 3C colored lines are representations of a global fit of a model with an 8-state sequential recognition to all data (See Supplementary Text and Fig. S5).

**Fig. S5.**
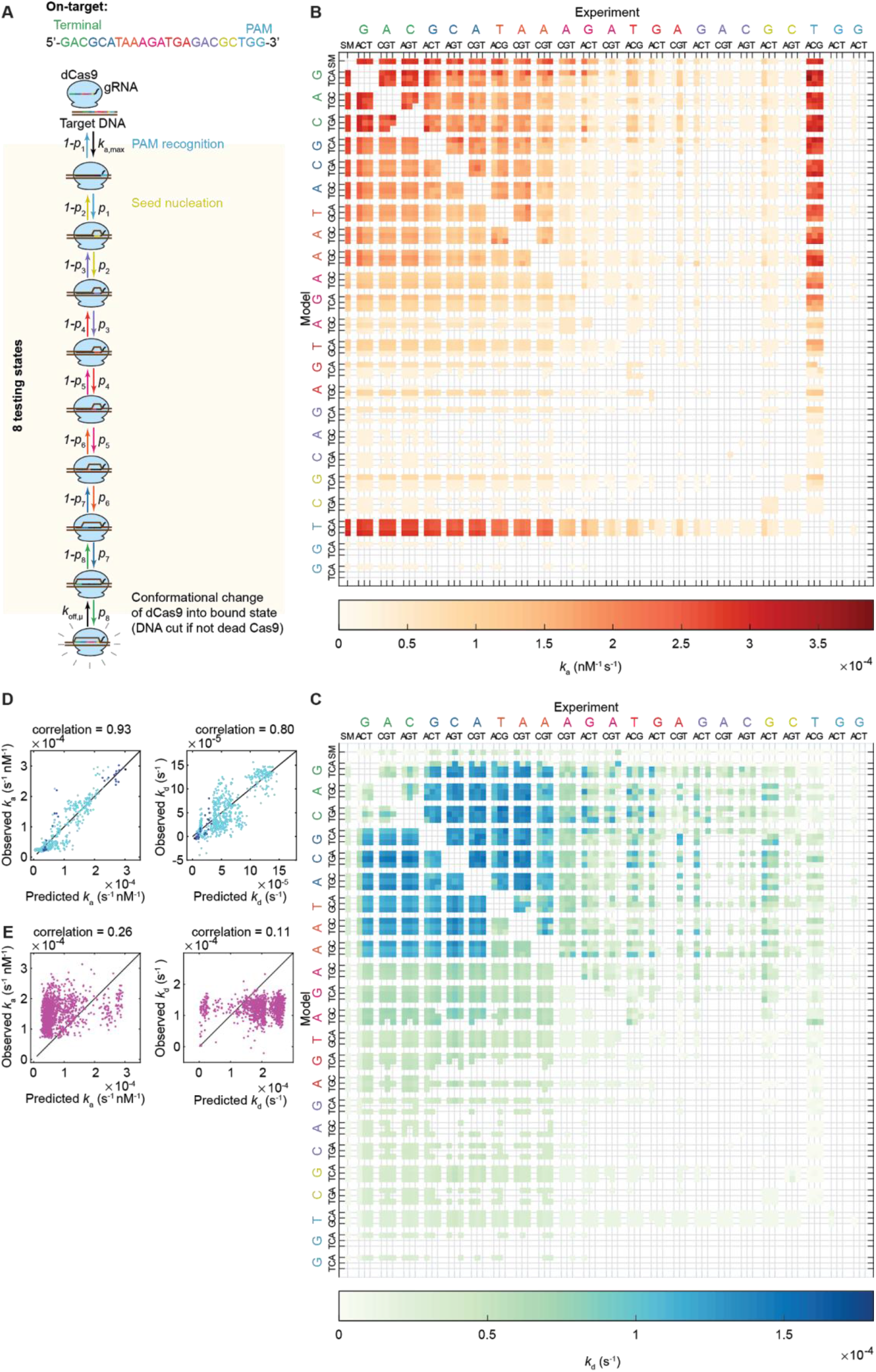
Model fit of dCas9 off-target binding and unbinding data. (**A**), Cartoon model diagram. Measured (above diagonal) and model-predicted (below diagonal) association (**B**) and dissociation (**C**) rates for single and double off-target mutants, when simultaneously fitting association and dissociation rates. The on-target, all single mutants, and all double mutants except the ones containing a mutation in the degenerate PAM (T in TGG) were used in the training data set. (**D**) Correlation between the model-predicted and observed association (left) and dissociation (right) rates when half of the data set (random sample) in (**B**,**C**) was used for training, and the other half of the data set was used for testing (points in the plots), for the on-target (black), single (blue) and double mutants (cyan). (**E**), Correlation between the predicted and observed association (left) and dissociation (right) for triple mutants when the same data set as in (**B**,**C**) is used for training the model.

### Supplementary Text

#### Derivation of Equation 1 and the coupling between macroscopic association and dissociation

The mean first passage times for transitions between the different states of a continuous time Markov chain are given by

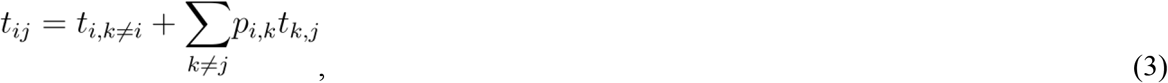

where *i*, *j* and *k* are state indices, *t*_i,j_ is the mean first passage time to transition from state *i* to state *j, t*_i,k≠i_ is the mean time to exit from state *i* into any of the other states in the model, and *p*_i,j_ is the probability to transition to state *j* given that the model is currently in state *i*.

We now consider the three-state model shown in Fig. 1A of the main text and evaluate Eq. 3 for all possible transitions in the model, which gives two linear systems of equations

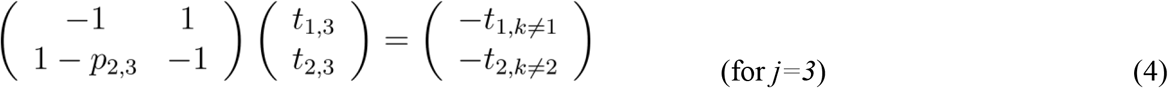

and

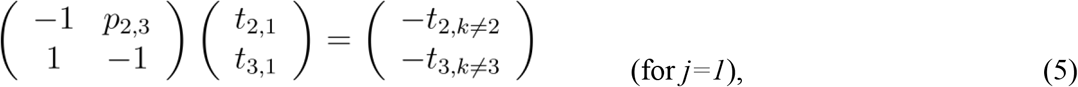

where we have used the fact that *p*_2,1_ = 1 -*p*_2,3_. We now solve Eq. 4 and 5 for 1/*t*_1,3_ and 1/*t*_3,1_, which are the sought macroscopic association and dissociation rates. We then obtain

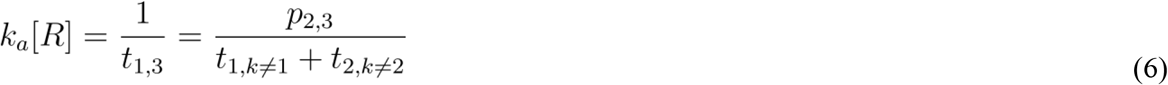

and

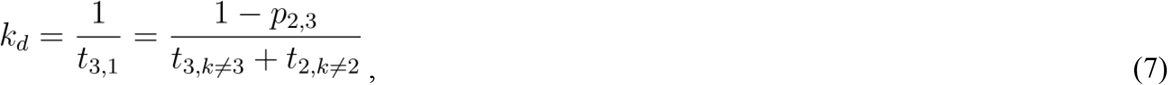

where [*R*] is the concentration of the searching protein. We now solve for *p*_2,3_ (= *p*_tot_) in both Eq. 6 and 7, and equate the resulting expressions, which gives

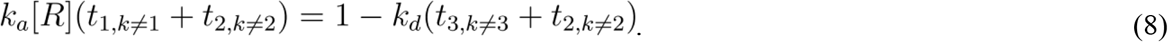

We then assume that *t*_2,k≠2_ ≪ *t*_1,k≠1_ and that *t*_2,k≠2_ ≪ *t*_3,k≠3_, i.e. that the time spent nonspecifically bound is much shorter than both the time spent specifically bound, and the time spent dissociated from the relevant DNA region, so that the *t*_2,k≠2_ terms in Eq. 8 are negligible. We now solve for *k*_a_[*R*], which gives

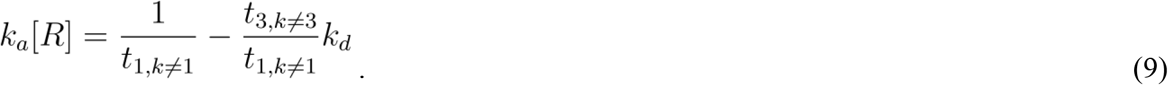

Using the same notations as in the main text, where *t*_1,k≠1_ = 1/*k*_on,max_[*R*] and *t*_3,k≠3_ = 1/*k*_off,μ_, yields the final expression,

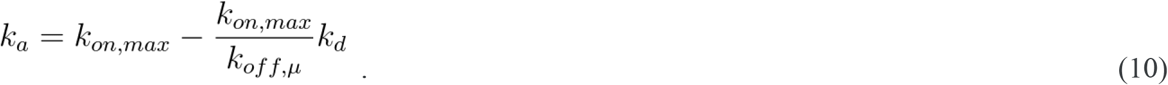

#### Extension of the model to handle multiple states of testing

The modeling framework can also be extended and used to determine the reaction mechanisms of binding and unbinding, if presented with measured kinetic rates of many binding site mutants. To achieve this, we extended the model to feature *N = n* +2 number of states, and *n* number of testing states that the protein has to proceed through sequentially to reach the specifically bound state, and where each testing state represents a group of nucleotides in the target sequence. The generalized model is thus

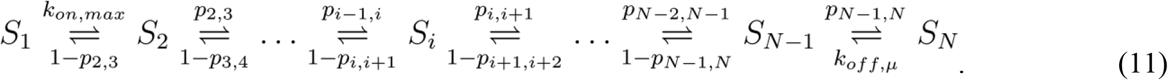

The model is parameterized by two rates *k*_on,max_ and *k*_off,μ_, defined by the diffusion-limited association time and the time spent specifically bound, along with *n* probabilities *p*_i,i+1_, defining how likely it is for the protein to transition from one testing state to the next state in the sequential binding. The two linear equation systems obtained after evaluating Eq. 3 for the state transitions in the model are now

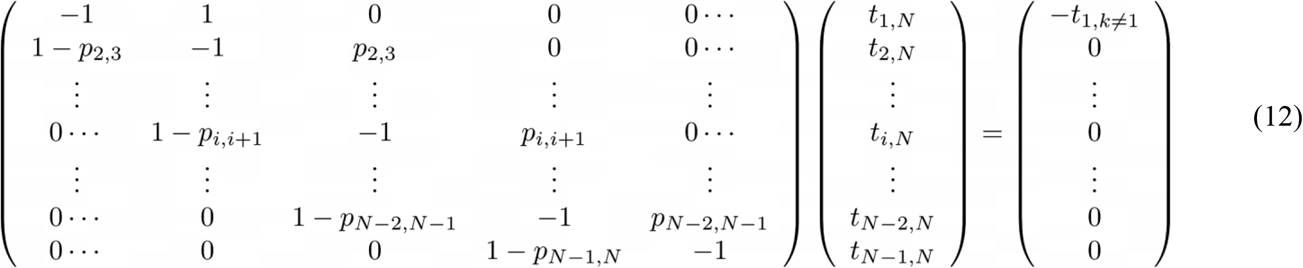

and

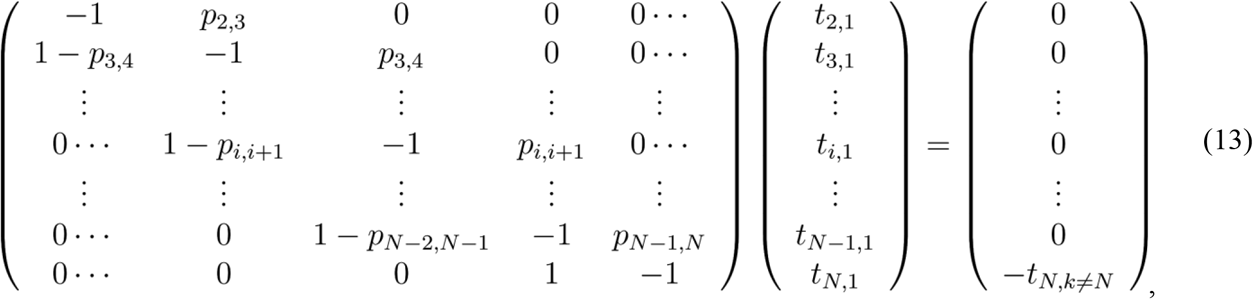

where we have used the assumption that the time spent in the nonspecific testing states is negligible, i.e. that *t*_i,k≠i_ = 0 for all *i* ∈ [2,*N*-1], and note that *t*_1,N_ = 1/*k*_a_[*R*], *t*_N,1_ = 1/*k*_d_, *t*_1,k≠1_ = 1/*k*_on,max_[*R*] and *t*_N,k≠N_ = 1/ *k*_off,μ_.We now consider how changes in one specific *p*_i,i+1_ parameter will change the *k*_a_ and *k*_d_ obtained from solving Eqs. 12 and 13, while keeping the rest of the model parameters constant. Since the time spent in any of the testing states is assumed to be 0, effectively the model is always in state 1 (free protein) or state *N* (bound protein). Hence, the passage times for reaching state *i* from state 1 must be exponentially distributed, with an average passage time of 1/*k*_on,max_*P*_1,i_, where *P*_1,i_ is the probability that the model will reach state *i* given that the model has just left state 1, before returning to state 1 again. Similarly, the passage times required to go from state *N* to state *i* must also be exponentially distributed, with an average passage time of 1/*k*_off,μ_*P*_N,i_, where *P*_N,i_ is the probability that the model will reach state *i* given that the model has just left state *N*, before returning to state *N* again. If we then view states {1, 2,…*i*-1} as one state, state *i* as one state, and states {*i*+1,*i*+2,…*N*} as one state, this new model describes a three-state continuous time Markov chain, which we have shown to have *k*_a_ and *k*_d_ coupled by Eq. 1. In this case, when considering the effect of changing one *p*_i,i+1_ parameter, we can rewrite Eq. 1 as

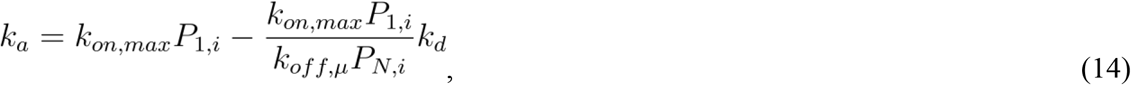

where *P*_1,i_ and *P*_N,i_ are functions of *p*_j,k_, where *j* ≠ *i* and *k* ≠ *i*+1. With *k*_on,max,i-1_ = *k*_on,max_*P*_1,i_ and *k*_off,μ,i-1_ = *k*_off,μ_*P*_N,i_, we obtain the same notation as used in Fig. 3, where *k*_off,μ,i-1_ is the effective rate of transitioning to state *i* from state *N*. Using this notation in Eq. 14 gives an equation of the familiar form

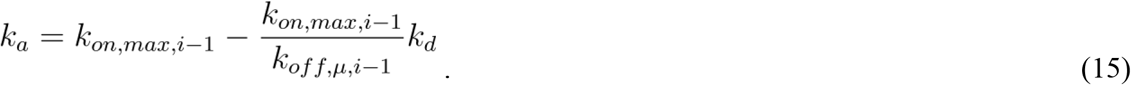

The model described by Eqs. 12 and 13 can be used to show effects of mutations in certain sequence regions (certain *p*_i,i+1_) on association and dissociation, which can be compared with experimental rates to deduce which nucleotides the protein contacts first, second, and so on in the recognition process of binding. To demonstrate this, we have presented such a model (Fig. S5A) with high-throughput data of association and dissociation available for dCas9 binding to off-target, mismatch mutants ((*14*), Fig. 3C). dCas9 is guided by an RNA (gRNA) when it binds DNA, with a reaction coordinate for the testing of recognition that is already well-established (*15*-*17*). Cas9 first detects a protospacer adjacent motif (PAM, NGG sequence), which is followed by DNA melting and gRNA-DNA hybridization, were the gRNA binds the DNA by base pairing in a sequential manner starting from a seed sequence, and then continuing hybridisation at base pairs more distal from the seed. If this sequential binding is completely memoryless, DNA binding sites with mismatches corresponding to gRNA region *j* should have the same microscopic dissociation rate (*k*_off,μ,j_) for the transition from a completely hybridized and specifically bound gRNA, to a melted region *j*. Just as our theory predicts according to Eq. 15, DNA binding sites with mutations in the same DNA region have association rates that are anti-correlated to the dissociation rates (Fig. 3C), where the *k*_d_-intercept of each *k*_a_ versus *k*_d_ line is the effective microscopic dissociation rate *k*_off,μ,j_ associated with each DNA region. When mismatches are present in the seed sequence, the slope of the *k*_a_ versus *k*_d_ line line is steep (low *k*_off,μ,j_ and mostly *k*_a_ modulation). The further away from the seed sequence the mismatches are, the flatter the slope becomes (higher *k*_off,μ,j_ and more *k*_d_ modulation).

We fit a model with eight-step sequential recognition to the data, that is with eight regions in the on-target DNA and one *p*_i,i+1_ parameter fitted for each unique sequence region, by solving Eq. 12 and 13 for 1/*t*_1,N_ and 1/*t*_N,1_ and choosing the parameters (*k*_on,max_,*k*_off,μ,n_, *p*_2,3_…) so that the summed squared deviation between the model predicted (*k*_a_,*k*_d_) and the experimentally obtained (*k*_a_,*k*_d_) is minimized. In total the model has 199 parameters, which are fitted to 2586 data values. With this model we then show the predicted effect of mutations for the individual DNA regions (colored lines in Fig. 3C, varying one *p*_i,i+1_ for each line). The model fit captures the large-scale changes and sequence-dependent coupling between *k*_a_ (Fig. S5B) and *k*_d_ (Fig. S5C) for single-and double-mismatch mutants, also when the model is trained with only half of the measured *k*_a_ and *k*_d_ values, while being tested on the other half of the dataset (Fig. S5D). We note that the model is simplistic since it assumes that the testing of recognition is infinitely fast. Since dCas9 is observed to bind more off-targets than Cas9 can cleave with high efficiency (*16*), a more realistic model would be one where the time that Cas9 actually spends in the testing state is considered (*25*). This discrepancy is a likely reason why our simple model is bad at predicting rates for triple mutants when trained on single and double mutants (Fig. S5E).

#### Measurements of LacI operator binding in relation to previous work

In one of our previous papers, we performed salt titrations with LacI labeled with bifunctional rhodamine (LacI-R) and detected the binding of *O*_1_ operators via single-molecule FRET (*8*).

From the titration end points of these measurements we obtained *K*_d_, *k*_a,obs_ and *k*_d,obs_ estimates of 0.0974 ± 0.0005 nM, 0.0033 ± 0.0003 s^−1^nM^−1^ and 0.0062 ± 0.0005 s^−1^ at 1 mM supplemented NaCl, or of 3.4 ± 0.6 nM, 0.0009 ± 0.0001 s^−1^nM^−1^ and 0.0089 ± 0.0007 s^−1^ at 80 mM supplemented NaCl. We note that these previous measurements, when compared with the measurements in this current work, were performed in a baseline imaging buffer with substantially higher ionic strength (10% glucose, 10% glycerol, 1mM NaCl, 0.05mM EDTA, 0.01% Tween 20, 0.1mg/ml BSA, 1mM 2-Mercaptoethanol, 2 mM Trolox, and 100mM K_2_HPO_4_:KH_2_PO_4_ pH 7.4 in the previous work versus 20mM K_2_HPO_4_:KH_2_PO_4_ pH 7.4 in this current work), which makes it most suitable to compare the results in this current work with measurements from the previous work that were obtained with ~100 mM lower concentrations of supplemented NaCl. The earlier estimates above should therefore be compared with the *K*_d_, *k*_a_, and *k*_d_ values for LacI-Far-2 at 100 mM or 200 mM supplemented NaCl in this work (0.025 ± 0.028 nM, 0.088 ± 0.029 s^−1^nM^−1^ and 0.0014 ± 0.0017 s^−1^ or 0.13 ± 0.07 nM, 0.023 ± 0.0096 s^−1^nM^−1^, and 0.0023 ± 0.0004 s^−1^, respectively). Furthermore, the previous work uses a LacI construct labeled at a different location, with which rates were measured using a different method. Most likely, the discrepancy between estimates from the current and earlier work is predominantly caused by the difference in how association events are detected. For the single-molecule FRET assay, more stringent selection occurred in the detection of association events, since the associating protein and the fluorophore must adopt a specific conformation capable of producing an interpretable FRET signal. In our single-molecule colocalization measurements, the only requirement for the detection of an association event is that the protein must contain an intact fluorophore label. Effectively, when normalizing the association rate to the concentration of labeled protein, single-molecule FRET is expected to yield lower apparent association rate constants compared to the single-molecule colocalization measurements, just as observed when comparing the measurements for LacI-R with those for LacI-Far-2. We note that our previous work focused on measuring intramolecular properties of target search when LacI scans the DNA via 1D diffusion, and that the absolute value of the apparent intermolecular association rate constant *k*_a_,obs reported there does not influence any of the conclusions. Prior to our work, the *K*_d_ for the full length LacI tetramer binding to a 80bp DNA fragment with the *O*_1_ operator has been estimated via nitrocellulose filter binding to be 0.16 nM at 50 mM KCl, and 0.43 nM at 100 mM KCl (*26*). Taken all together, we believe that the discrepancy in *K*_d_ estimates is in line with what can be expected from measurements with different protein constructs and batches that were carried out in different buffers and with different experimental methods.

If we assume that the observed *k*_d_,obs at 1 mM NaCl in the previous work is predominantly due to photobleaching (*k*_d_= 0 and *k*_a_ = *k*_on,max_ for this salt concentration), we can obtain estimates of *k*_d_ for the other salt concentrations in the single-molecule FRET measurements by fitting a linear equation (Eq. 1) to the six salt concentration data points, and by subtracting the *k*_d_ value corresponding to *k*_a_ = *k*_on,max_ from all the data points. The *k*_d_-intercept of this photobleaching-corrected line is then *k*_off,μ_. When we perform this analysis, we obtain *k*_off,μ_ = 0.0031 ± 0.0002 s^−1^, which is identical to the *k*_off,μ_ value for LacI-Far-2 binding to *O*_1_ as obtained in our single-molecule colocalization measurements in Fig 1H. For LacI-R, error bars in this section are standard errors obtained by propagating and resampling the experimental errors, while assuming that the errors reported in (*8*) are normally distributed, and the *K*_d_ estimates are reported after correcting for photobleaching as described above. For LacI-Far-2, error bars are standard errors with *n* = 2 independent experiments.

#### Energy landscape for reaching the specific bound state via the putative reaction coordinate

To estimate and draw the energy landscapes *in vivo* for the different operators in Fig. 2D, we first consider the free energy difference between the free (state 1) and bound (state 3) state. Since the time spent in the testing state is much shorter than the time spent in the free and bound states, this free energy difference is directly given by the measured *K*_d_ values according to

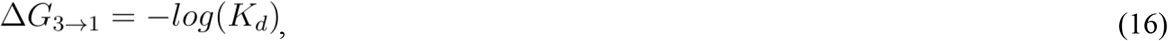

with the energy difference given in *k*_b_T units. Next, we consider the free energy difference between the free state and the testing state, which is the same for all operators. This free energy difference is defined as

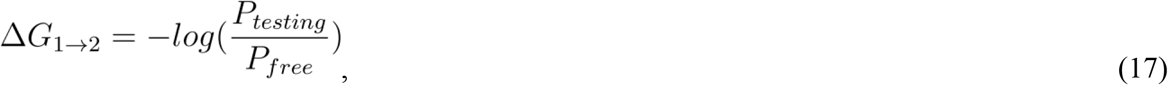

where *P*_testing_ is the probability that the protein is testing for recognition within one sliding length from the operator, and *P*_free_ is the probability that the protein is searching somewhere else in the cell at any given time point. This probability can be calculated according to

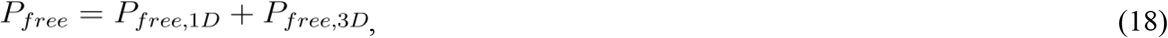

where *P*_free,1D_ is the probability that the protein is bound nonspecifically to DNA somewhere else in the genome of the cell, and *P*_free,3D_ is the probability that the protein is dissociated from DNA and is searching in the cytoplasm. The fraction of time *f* that the protein spends nonspecifically bound to DNA when searching is thus defined as

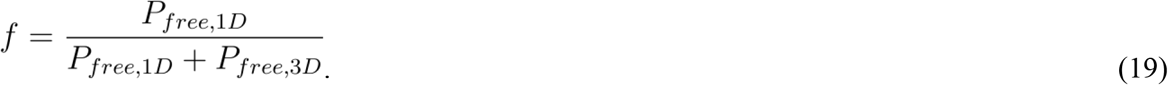

After combining Eqs. 17–19 we obtain

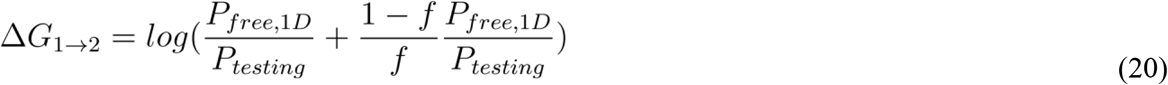

where the ratio *P*_free,1D_ /*P*_testing_ can be calculated from the size of the genome *N*_genome_ and the average sliding length *N*_sliding_ as

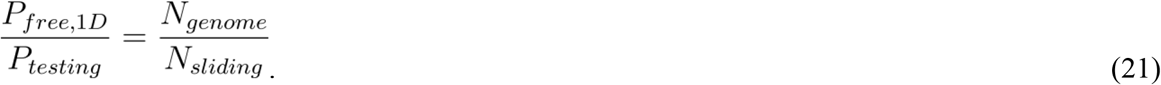

With *N*_genome_ = 5×10^6^ base pairs, *N*_sliding_ = 45 base pairs (*7*), and *f* = 0.9 (*27*) we obtain the free energy difference shown in Fig. 2D.

The relative difference in activation energy on the transition path between the testing state (state 2) and bound state (state 3) for the different operators can be calculated from the measured *k*_off,μ_ values. To achieve this, we model the transition state that the protein has to go through to reach the bound state from the testing state as a distinct species. This modeling scheme is similar to what is used in transition state theory (TST), but here we model the transition state as a true equilibrated Markovian state, instead of as a quasi-equilibrated state as is done in TST. The model is thus

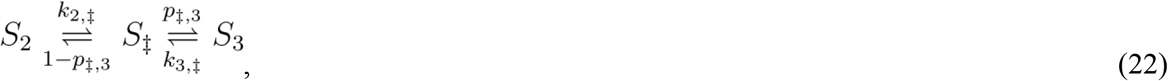

where *S*_2_ is the testing state, *S*_‡_ is the transition state, and *S*_3_ is the bound state. As we have shown, Eq. 7 describes how the effective transition rate from the last state to the first state in this type of model can be calculated. This effective rate is now *k*_off,μ_, and with the notations used in Eq. 22, Eq. 7 is for this model

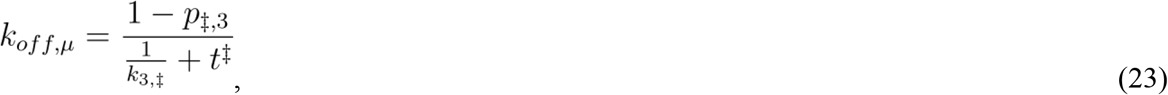

where *t^‡^* is the average time that the model spends in the transition state. Furthermore, the free energy difference between the bound state and the transition state is defined as

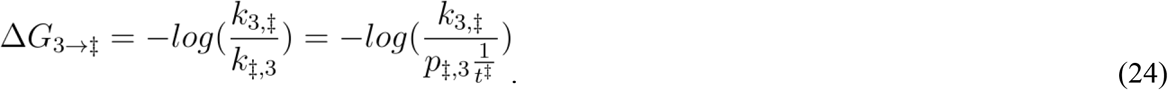

Solving for *k*_3,‡_ in Eq. 23 and putting this into Eq. 24 gives

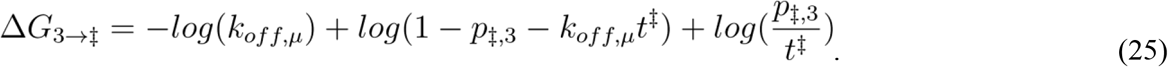

When 1/*k*_off,μ_ >> *t*^‡^, i.e. when the time spent in the transition state is very small, Eq. 25 can be written as

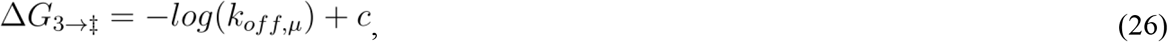

where

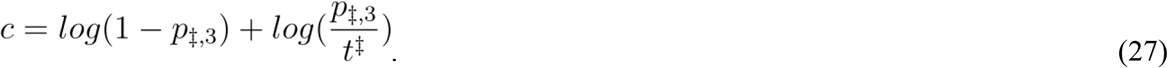

If we now assume that *t*^‡^ and *p*_‡,3_ are the same for all operators, the difference in Δ*G*_3→‡_ for different operators, i.e. the difference in energy barrier between the bound state and the transition state for different operators in Fig. 2D, is given by the difference in −log(*k*_off,μ_) for the different operators, where a fast *k*_off,μ_ gives a low energy barrier, and a slow *k*_off,μ_ gives a high energy barrier.

**Table S1.**
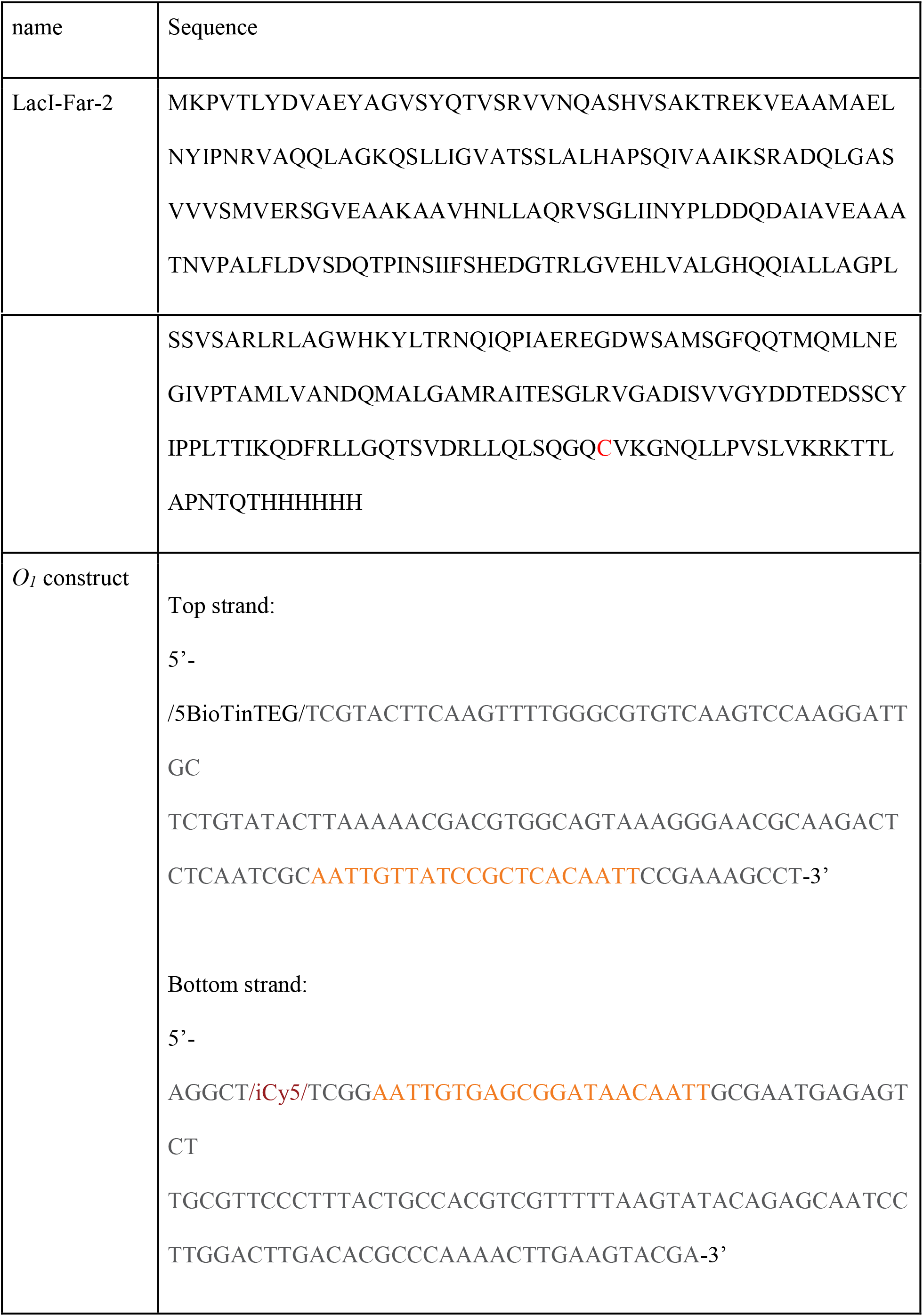
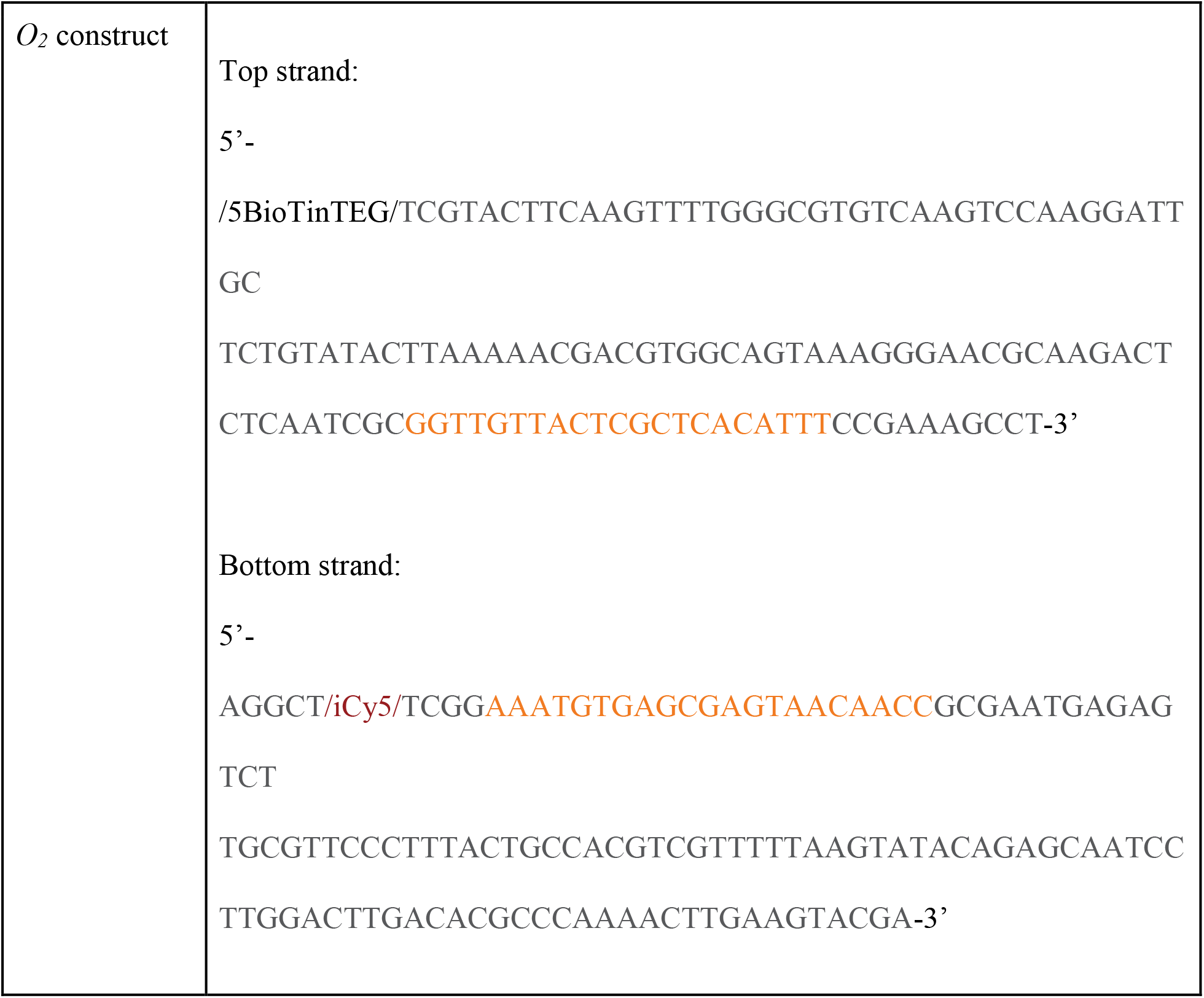
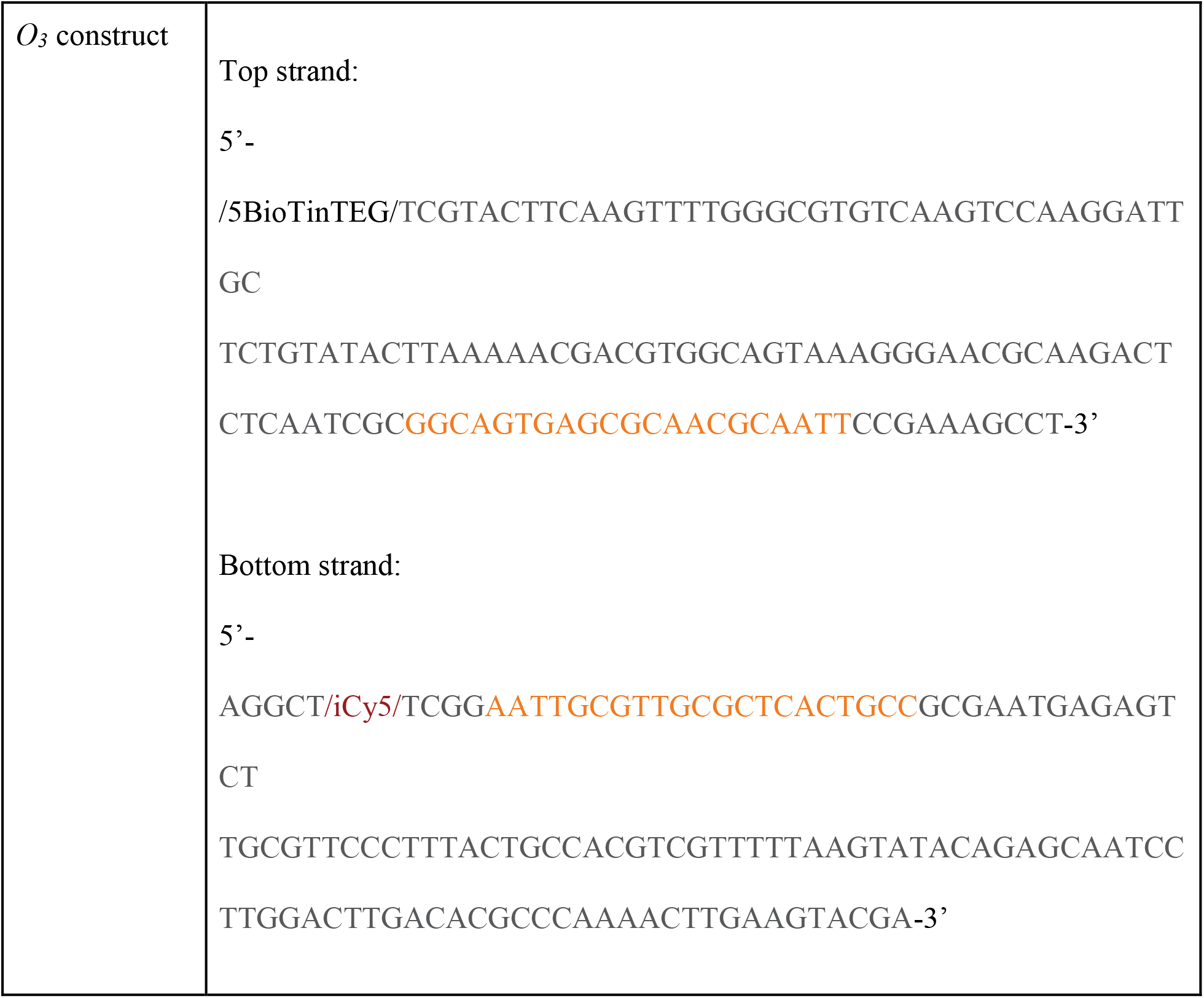
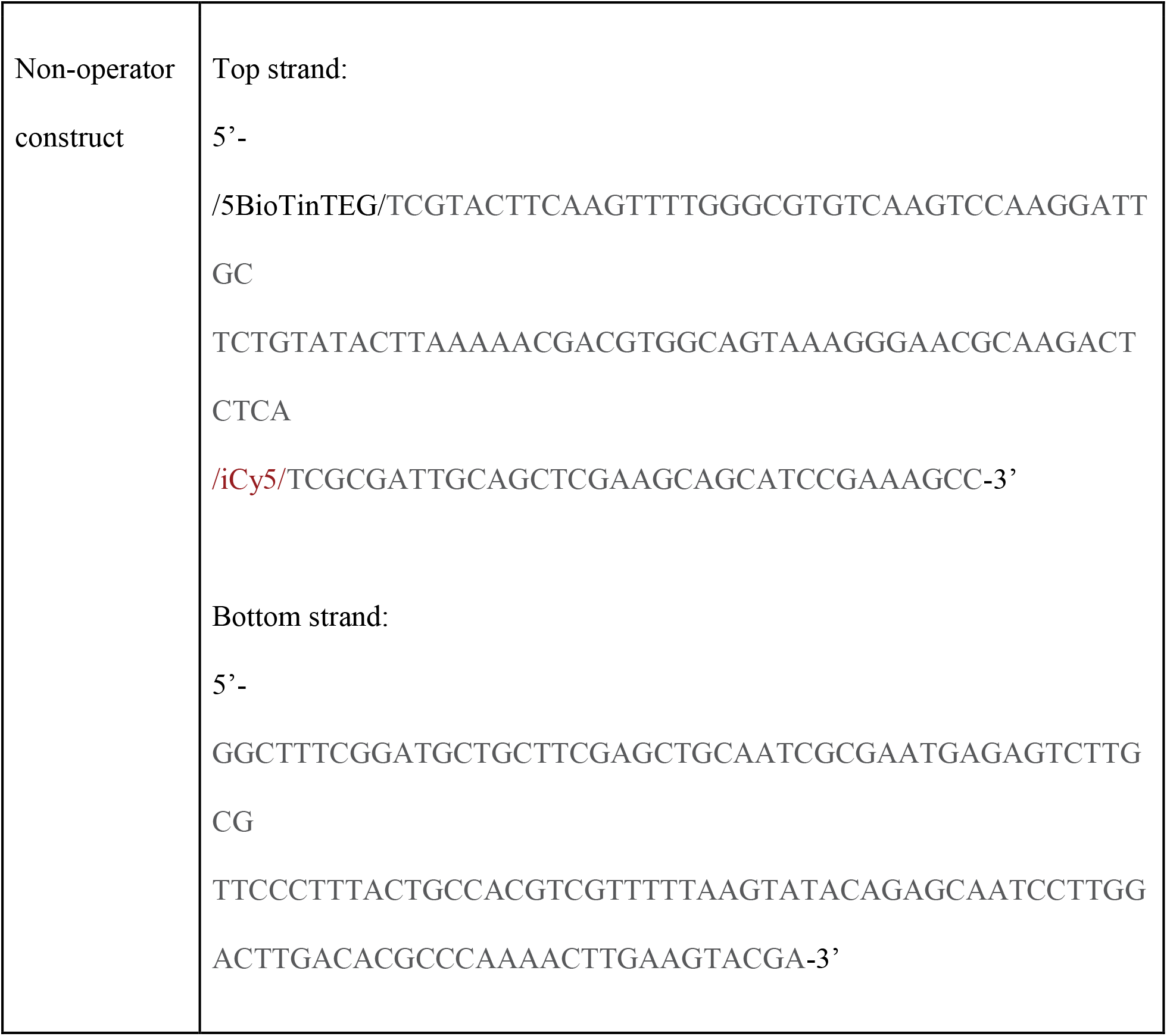
Sequence of LacI and DNA constructs for single-molecule colocalization measurements. For LacI-Far-2, the cysteine introduced for labeling is marked in red. For the DNA constructs, modifications are reported using Integrated DNA Technologies (IDT) nomenclature. LacI operator sites are highlighted in orange.

**Table S2.** Stochastic simulations and simulated microscopy. True values of *k*_off,μ_ for *k*_a_ and *k*_d_ values and lines put into stochastic simulations, and estimated *k*_off,μ_ values obtained after running simulated microscopy and the analysis pipeline. Estimations of *k*_off,μ_ are given as 68% confidence intervals obtained by bootstrapping the simulated fluorescence traces. See Fig. S4 for true and estimated *k*_a_ and *k*_d_ values for these *k*_off,μ_ lines.

|  |  |  |  |  |  |  |
| --- | --- | --- | --- | --- | --- | --- |
| True $k_{\text{off},\mu}$<br>($\text{s}^{-1}$) | $1 \cdot 10^{-4}$ | $1 \cdot 10^{-3}$ | $1 \cdot 10^{-2}$ | $2 \cdot 10^{-2}$ | $3 \cdot 10^{-2}$ | 100 |
| Estimated<br>$k_{\text{off},\mu}$ ($\text{s}^{-1}$) | [-3.6,1.1]<br>$\cdot 10^{-4}$ | [0.84,1.2]<br>$\cdot 10^{-3}$ | [0.89,0.96]<br>$\cdot 10^{-2}$ | [1.8,1.9]<br>$\cdot 10^{-2}$ | [2.5,2.7]<br>$\cdot 10^{-2}$ | [8.7,∞]<br>$\cdot 10^{-2}$ |

